# Three previously characterized resistances to yellow rust are encoded by a single locus *Wtk1*

**DOI:** 10.1101/2020.01.02.892968

**Authors:** Valentyna Klymiuk, Andrii Fatiukha, Dina Raats, Valeria Bocharova, Lin Huang, Lihua Feng, Samidha Jaiwar, Curtis Pozniak, Gitta Coaker, Jorge Dubcovsky, Tzion Fahima

## Abstract

The wild emmer wheat (*Triticum turgidum* ssp. *dicoccoides*; WEW) yellow (stripe) rust resistance genes *Yr15, YrG303* and *YrH52* were discovered in natural populations from different geographic locations. They all localize to chromosome 1B but were thought to be non-allelic based on differences in resistance response. We recently cloned *Yr15* as a *Wheat Tandem Kinase 1* (*WTK1*) and showed here that these three resistance loci co-segregate in fine-mapping populations and share identical full-length genomic sequence of functional *Wtk1*. Independent EMS mutagenized susceptible *yrG303* and *yrH52* lines carried single nucleotide mutations in *Wtk1* that disrupted function. A comparison of the mutations for *yr15, yrG303* and *yrH52* mutants showed that while key conserved residues were intact, other conserved regions in critical kinase subdomains were frequently affected. Thus, we concluded that *Yr15-, YrG303-* and *YrH52*-mediated resistances to yellow rust are encoded by a single locus *Wtk1*. Introgression of *Wtk1* into multiple genetic backgrounds resulted in variable phenotypic responses, confirming that *Wtk1*-mediated resistance is part of a complex immune response network. WEW natural populations subjected to natural selection and adaptation have potential to serve as a good source for evolutionary studies of different traits and multifaceted gene networks.

**Highlight:** We demonstrate that *Yr15, YrG303* and *YrH52* resistances are encoded by the *Wtk1* locus, but express variable resistance responses to yellow rust in a genetic background dependent manner.

## Introduction

Wheat has been the basic staple food for the major civilizations of Europe, West Asia and North Africa for at least 10,000 years (Nevo *et al.*, 2002). Today, common wheat (*Triticum aestivum* L.) and durum wheat (*T. turgidum* ssp. *durum* (Desf.) Husnot) provide 20% of the calories and proteins for human consumption, as well as vitamins, dietary fibers, and phytochemicals (Shewry and Hey, 2015). Wheat annual yield reaches more than 900 million tons (Food and Agriculture Organization Corporate Statistical Database, FAOSTAT); however, losses due to biotic (pathogens) and abiotic (unfavorable growth conditions) stresses prevent the maximum yield potential from being achieved.

Wheat yellow rust, also known as a stripe rust, is caused by the basidiomycete fungus *Puccinia striiformis* f. sp. *tritici* (*Pst*), an obligate pathogen that threatens wheat production around the globe (Chen, 2005). Yield losses due to yellow rust have ranged from 10% to 70% in susceptible varieties and a total yield loss (100%) can occur under severe epidemics (Chen, 2005). Host resistance is considered to be the most economically and environmentally friendly strategy for yellow rust control (Chen, 2005), but widespread use of initially effective resistance genes can lead to rapid breakdown of resistance, e.g. the appearance of *Pst* races that overcome widely deployed R-genes, such as *Yr2, Yr9, Yr17*, and *Yr27*, has led to destructive pandemics (Wellings, 2011; Hovmøller *et al.*, 2015). Moreover, the rapid evolution of the pathogen facilitates an expansion of *Pst* into new regions, and therefore becoming an emerging issue in some countries, such as western Canada (Brar *et al.*, 2018). Thus, resistant wheat variety breeding is a continuous process to withstand yellow rust epidemics globally, using all possible sources of *Pst* resistance, in order to widen and diversify the existing R-gene pool (Roelfes *et al.*, 1992).

Wild emmer wheat (WEW), *T. turgidum* ssp. *dicoccoides* (Körn. ex Asch. & Graebner) Thell. (BBAA), discovered in 1906 in Rosh Pina, Israel by A. Aaronsohn (Aaronsohn, 1910), has been recognized as an important source for novel yellow rust resistance (*Yr*) genes (Fahima *et al.*,1998; Huang *et al.*, 2016a; Klymiuk *et al.*, 2019a). WEW is the undomesticated polyploid progenitor for modern cultivated tetraploid durum wheat (BBAA) and hexaploid common wheat (BBAADD) and natural populations still grow in a wide range of ecogeographical conditions distributed across the Near East Fertile Crescent (Özkan *et al.*, 2011). These natural populations can serve as a model to study the evolutionary processes that shaped the currently observed allelic variation (Yahiaoui *et al.*, 2009; Sela *et al.*, 2011; Huang *et al.*, 2016b; Lundström *et al.*, 2017; Klymiuk *et al.*, 2019b). Several previous studies have reported that WEW accessions exhibit high levels of resistance to inoculation with *Pst* isolates (Gerechter-Amitai and Stubbs, 1970; Van Silfhout, 1989; Nevo *et al.*, 1986). Initially, *Yr* genes were considered novel if they expressed distinct reaction patterns to a set of *Pst* isolates. With this definition, Van Silfhout (1989) predicted the presence of at least 11 major *Yr* genes in WEW populations. Currently, the genetic position of an identified gene compared with these of previously mapped loci is considered to determine novelty (McIntosh *et al.*, 2017). So far, six WEW*-*derived *Yr* genes have been recognized (*Yr15, YrH52, Yr35, Yr36, YrTz2*, and *YrSM139-1B*) on chromosome arms 1BS and 6BS (Huang *et al.*, 2016a; Klymiuk *et al.*, 2019a). Among these, only *Yr36* (Fu *et al.*, 2009) and *Yr15* (Klymiuk *et al.*, 2018) have been cloned so far.

*Yr15* was discovered in WEW accession G25 (G25) by Gerechter-Amitai *et al.* (1989) and was shown to confer resistance against a worldwide collection of more than 3000 genetically diverse *Pst* isolates (Sharma-Poudyal *et al.*, 2013; Ali *et al.*, 2017; Liu *et al.*, 2017; Chen and Kang, 2017). Only a few isolates virulent on *Yr15* from Afghanistan (Van Silfout 1989) and Denmark (Hovmøller and Justesen, 2007) have been reported. *Yr15* was localized to the short arm of chromosome 1B (Sun *et al.*, 1997; Peng *et al.*, 2000b; Yaniv *et al.*, 2015), and its positional cloning revealed that it encodes for a protein with 665 amino acid residues (Klymiuk *et al.*, 2018). Yr15 protein was designated as Wheat Tandem Kinase 1 (WTK1) since it possesses a Tandem Kinase-Pseudokinase (TKP) domain architecture and phylogenetically groups with other proteins sharing similar TKP structure (Klymiuk *et al.*, 2018). Functional (*Wtk1*) and non-functional (*wtk1*) *WTK1* alleles differ by insertions of transposable elements, indels and stop codons (Klymiuk *et al.*, 2018). Functional markers detected alternate alleles in WEW, *T. turgidum* ssp. *durum*, and *T. aestivum* that were consistent with the phenotypic responses (Klymiuk *et al.*, 2019b).

*YrH52* was also identified in WEW and its introgression into adapted cultivars of durum wheat and bread wheat provides effective resistance to *Pst* (Peng *et al.*, 1999; Klymiuk *et al.*, 2019a). A primary genetic map of *YrH52* on chromosome 1BS was developed by Peng *et al.* (1999,2000a,b) using a segregating population from a cross between WEW accession H52 (the donor line of *YrH52*) and *T. turgidum* ssp. *durum* cv. Langdon. *YrG303* was identified in the WEW donor line G303 and was shown to provide resistance to 28 *Pst* isolates from 19 countries tested by Van Silfhout (1989).

*Yr15, YrG303*, and *YrH52* all localize to the distal region of chromosome 1B of wheat. In the current study, we used positional cloning and functional validation to demonstrate that *Yr15, YrG303*, and *YrH52* are encoded by the functional *Wtk1* allele. Nevertheless, WEW accessions and introgression lines that carry *Yr15, YrG303*, and *YrH52*, show different phenotypic responses after challenged with *Pst*. Mutations in WTK1 in the *yr15, yrG303* and *yrH52* susceptible mutants demonstrate conserved regions in critical kinase subdomains likely important for functionality.

## Materials and methods

### Development of mapping populations and introgression lines

#### YrH52

The *YrH52* mapping population used in the current study consisted of 3,549 F_2_ plants derived from a cross between resistant WEW (male) accession H52 (TD010027 from ICGB, Institute of Evolution, University of Haifa) collected in Mt. Hermon (N33°17’19” E35°45’18”), the donor of *YrH52* (**Fig. 1**), and a susceptible durum wheat (female) cv. Langdon (Peng *et al.*, 1999). A homozygous resistant BC_3_F_2_ Ariel-YrH52 introgression line (LDN/H52//3*Ariel) was used to develop EMS *yrH52* mutants.

**Figure 1.**
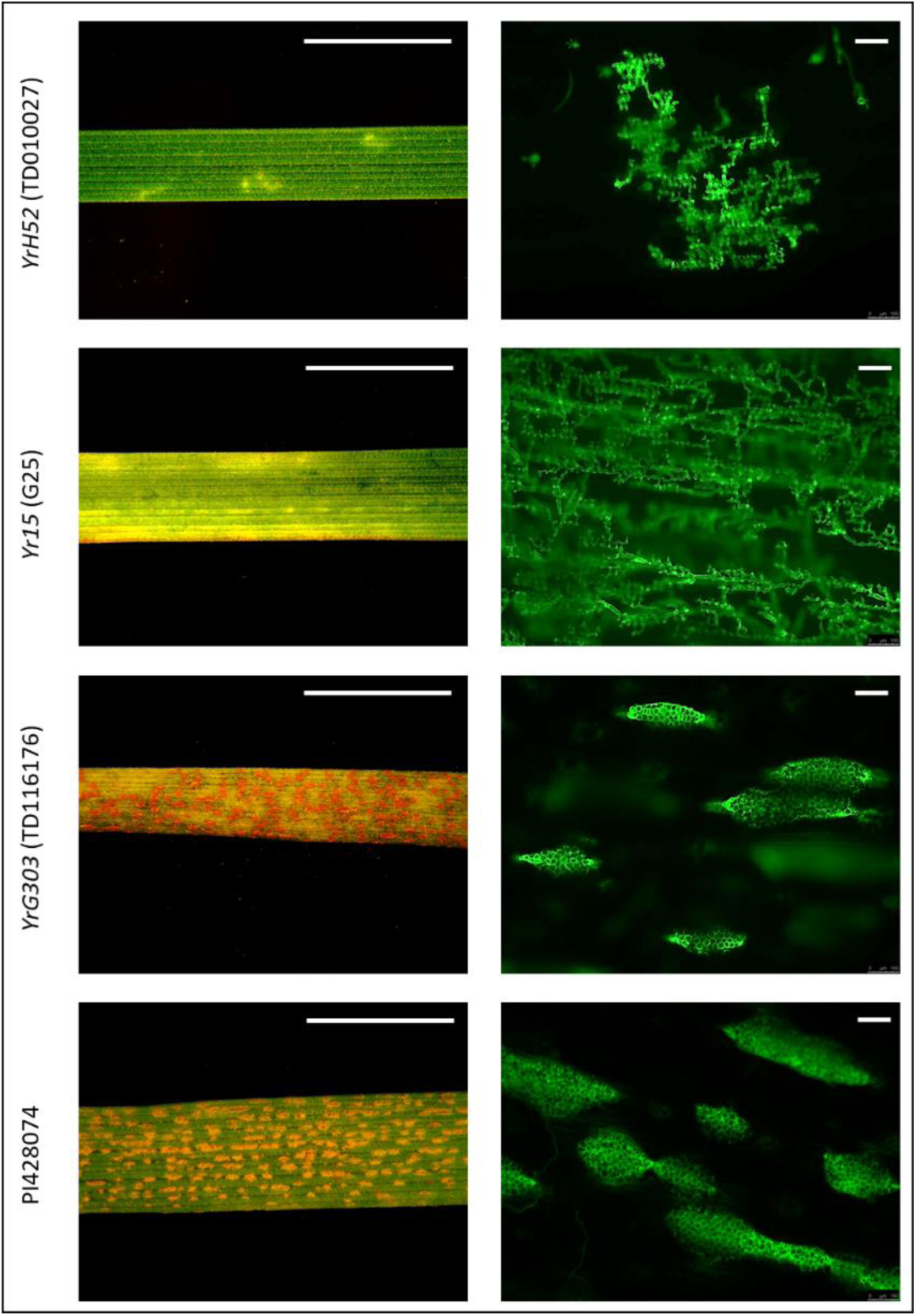
Differences in seedling phenotypic responses of WEW accessions donors of *YrH52, Yr15* and *YrG303*, and susceptible control PI428074 accession, at 14 dpi with *Pst* isolate #5006. Left panel shows binocular microscopic observation of hypersensitive response and fungal sporulation (Scale bar = 1 cm). Right panel represents sizes of fungal colonies and uredinia bags visualized by fluorescent dye WGA-FITC (Scale bar = 100 µm). Note that G303 plants exhibited variable levels of resistance responses upon inoculation at the seedling stage (IT1-5) and only predominant IT5 is presented here. This figure is available in colour at JXB online.

#### YrG303

An F_2_ population (1,917 plants) was developed for fine mapping of *YrG303* by crossing resistant WEW (male) accession G303 (TD116176 from ICGB, Institute of Evolution, University of Haifa) collected in Dishon, Israel (N33°05’09” E35°31’03”; Fahima *et al.*, 1998), the donor of *YrG303* gene (**Fig. 1**), with a susceptible *T. turgidum* ssp. *durum* accession D447 (LD393/2*Langdon ND58-322). A resistant hexaploid line 2298 (Vaskar*3/G303//Avocet) and resistant RILs of A95 mapping population derived from crossing 2298 with susceptible 2463 (Vaskar*3/G303//Avocet) were used for development of *yrG303* mutants.

All parental lines were confirmed to be near-homozygous using the 15K wheat SNP array (Muqaddasi *et al.*, 2017) (Trait Genetics GmbH, Gatersleben, Germany) consisting of 12,905 SNPs selected from the wheat 90 K array (Wang *et al.*, 2014).

### Yellow rust test under controlled growth chamber conditions

Yellow rust test was carried out with *Pst* isolate #5006 (race 38E134) using a standard protocol for *Pst* inoculation described in Klymiuk *et al.* (2019b). Ten plants per genotype were inoculated at seedling stage (the two-leaf stage). The yellow rust response variation was evaluated 14 to 18 days post inoculation (dpi) using a 0 to 9 scale of IT (Line and Qayoum, 1992) with the following interpretation of the results: IT = 0–3 were considered as resistant, IT = 4–6 as moderately resistant, IT = 7–9 as susceptible.

### Field experiment for yellow rust assessment at adult stage

A field experiment was conducted in order to evaluate *Pst* resistance of F_4_ recombinant lines at the adult stage. Homozygous recombinant lines from the tetraploid *YrG303* mapping population, related parental lines G303, D447, 2298, 2463 as well as susceptible control *T. aestivum* cv. Morocco were planted in the Institute of Evolution field located on Mt. Carmel (32°45’30.276’’ N, 35°1’21.7596’’ E) during the 2016-2017 winter growing season. Field design included three replicates per genotype randomly distributed within three 15 m long rows. Each replicate included three plants within a row planted at the distance of 30 cm between lines and 15 cm between plants. Seeds were germinated in rolled wet paper, kept in dark cold room (4°C) for three days, and then transferred to growth room (20°C) with 14/10 hours light/dark regime for 24h before sowing in soil. Only seeds that showed good germination and root development were transferred to the field. Plants were inoculated at flag leave stage (adult inoculation) using a sprayer with fresh *Pst* isolate #5006 spores suspended in Soltrol® 170 light oil (Chevron Phillips Chemical Company, The Woodlands, TX, USA) one hour before sunset at average day/night temperatures of 22/12°C. To ensure inoculation, spreaders inoculated in controlled chamber conditions were evenly distributed in pots in the field, every second line within rows. Phenotypes were scored 14, 21 and 29 dpi for each plant; the most advanced IT was considered as the final score.

### Histopathological characterization of Pst-wheat interactions within leaf tissues

Fluorescent visualization of *P. striiformis* structures during infection was conducted according to the protocol described by Dawson *et al.* (2015), with slight modifications. This protocol used wheat germ agglutinin (WGA; a lectin that binds specifically to β (1→4)-N-acetyl-D-glucosamine, i.e., chitin) conjugated with a fluorescent dye to visualize the intercellular fungal growth and pustule formation on infected leaves. Leaf segments (10 cm long, 2nd leaf) were sampled at 14 dpi from WEW accessions G25, G303 and H52 inoculated with urediniospores of *Pst* isolate #5006 as described above.

The sampled leaf segments were placed in 15 ml centrifuge tubes containing 15 ml of 1 M KOH and 2-3 drops of the surfactant alkylaryl polyether alcohol (Spreader DX) for clearing the tissue. Tubes were kept at 37°C for 24 h, followed by three washes of samples with 50 mM Tris (pH 7.5) for neutralization of pH. After the last wash, the leaves were gently transferred to a 9 cm plastic petri dishes, excess of Tris solution was removed and 1 ml of 20 µg/ml WGA conjugated to fluorophore Alexa 488 (L4895-2MG; Sigma-Aldrich) in 50 mM Tris was placed on the leaf surface. The leaves were stained with WGA for 24 h at 4°C, and then washed with ddH_2_O. Stained leaf tissues were gently placed on microscope slides, immersed with antifade mounting medium for preserving fluorescence (Vectashield, Vector Laboratories), covered with cover glass, sealed with rubber cement, and stored at 4°C in the dark. Fluorescence microscopy was performed on an inverted fluorescence microscope, Leica DMi8 (Leica Microsystems, Wetzlar, Germany), fitted with a filter cube for the FITC excitation range (Ex: 460–500; Dc: 505; Em: 512–542), and a FLUO regime to observe the WGA-stained fungal structures. Three plants of each wheat genotype were used for the investigation and whole 10 cm long leaf segments were examined in each case. Images of the most predominant fungal colonies/fields of view were recorded for each genotype.

### Development of CAPS and KASP markers

Detailed protocols for the development and use of cleaved amplified polymorphic sequences (CAPS) markers for the screening of mapping populations are described in Raats *et al.* (2014). Sequences of the newly developed CAPS marker *uhw290* is presented in **Table S1**.

For development of new KASP markers, the G303, D447, H52, and Langdon parental lines of the *YrG303* and *YrH52* mapping populations were genotyped using the 15K wheat SNP array. Polymorphic markers, residing on chromosome 1BS between *Yr15* flanking SSR markers *barc8* and *gwm273*, were identified based on their location on the consensus tetraploid wheat genetic map (Maccaferri *et al.* 2015). Sequences of eight single nucleotide polymorphism (SNP) markers (*RFL_Contig2160_617, IACX502, Ra_c16879_977, BS00087784_51, Excalibur_c17202_1833, wsnp_Ku_c4911_8795151, wsnp_Ex_c2111_3963161*, and *RAC875_c79370_378*) were converted into KASP markers using the software Polymarker (Ramirez-Gonzalez *et al.* 2015). The primer sequences of these KASP markers are presented in **Table S2**.

Detailed descriptions of the development, sequences and conditions for amplification of KASP *Yr15* functional molecular markers are provided in Klymiuk *et al.* (2019b). Names, primer sequences, and references of other previously published markers are presented in **Table S1**.

### Construction of genetic linkage maps

Primary genetic maps were developed for each of the genes, *Yr15* (Klymiuk *et al.*, 2018), *YrG303* and *YrH52*, using the following markers: SSR markers *barc8* and *uhw273*; KASP markers *RAC875_c826_839* and *BS00022902_51*; CAPS markers *uhw259* and *uhw264*. The primary genetic maps were constructed using the MultiPoint package (http://www.multiqtl.com/). For high-resolution mapping F_2_ plants from large mapping populations were screened first with two markers flanking each of the target genes. Plants that showed recombination events between the two markers were self-pollinated and progressed to F_3_. Ten to sixteen F_3_ plants of each of the F_2_ recombinants were analyzed with markers and homozygous RILs were selected. The F_3_ RILs were then screened with molecular markers that resided within the defined *YrG303* or *YrH52* interval. Phenotyping of F_3-4_ RILs for response to *Pst* inoculation was performed as described above. High-resolution genetic maps were constructed using the graphical genotypes approach (Young and Tanksley, 1989). Genetic distances obtained for the low-resolution maps, were used as a reference to estimate the relative genetic distances within the high-resolution maps.

### Collinearity between genetic and physical maps

The best hits of BLASTN search of the corresponding primer sequences for each genetic marker against the three genome assemblies of wheat, *T. dicoccoides* Zavitan (Avni *et al.*, 2017), *T. turgidum* ssp. *durum* Svevo (Maccaferri *et al.*, 2019), and *T. aestivum* Chinese Spring (Appels *et al.*, 2018), were used to estimate average physical distances between markers. Visualization of the genetic and physical maps was performed with MapChart 2.2 software (Voorrips, 2002).

### EMS mutagenesis and screening for susceptible mutants

Seeds of the homozygous resistant *T. aestivum* line 2298 carrying an introgression from WEW G303 that harbors the *YrG303* locus and 14 homozygous resistant F_6_ RILs of A95 population were treated with 0.4% ethyl methanesulfonate (EMS) following the protocol described in Klymiuk *et al.* (2018), in order to obtain susceptible *yrG303* mutants. EMS-treated M_1_ plants were grown in the Institute of Evolution field located on Mt. Carmel. Seedlings of M_2_ families of F_6_ RILs of A95 population (10-20 seeds per family) were artificially inoculated with *Pst* under field conditions as described above. While seedlings of M_2_ families of 2298 line (12 seeds per family) were screened under growth chamber conditions as described above.

Following the same EMS mutagenesis protocol (Klymiuk *et al.*, 2018), seeds of *T. aestivum* line Ariel-YrH52 that carry an introgression from WEW H52 line with *YrH52* locus were treated with 0.5% EMS for development of loss-of-function *yrH52* mutants. EMS-treated M_1_ plants were grown and seedlings of M_2_ families (10-20 seeds per family) were screened for the response to inoculation with *Pst* under field conditions at the Institute of Evolution field located on Mt. Carmel as described above.

All M_3_ families (*yrG303* and *yrH52*) obtained from M_2_ susceptible plants were inoculated in a growth chamber, as described above, to confirm homozygosity of the recessive mutations.

### Sequencing of WTK1 from the mutants

DNA was isolated from freeze-dried leaves of M_3_ plants using standard CTAB protocol (Doyle, 1991). Coding regions of *WTK1* were sequenced from each *yrG303* and *yrH52* mutant using WJKDF1/WJKDR1, WJKDF2/WJKDR2 and WJKDF3/WJKDR3 primer pairs (Klymiuk *et al.*, 2018) and screened for mutations. In order to confirm the absence of the detected mutations in the resistant background we sequenced *WTK1* regions spanning identified mutations from two M_2_-derived resistant sister lines for each of the *yrG303* mutant.

### Structure of WTK1 protein domains

Full-length sequences of *Wtk1* from G25, G303 and H52 were previously published (Klymiuk *et al.*, 2018; Klymiuk *et al.*, 2019b) and have been deposited in NCBI GenBank under accession numbers MG649384, MK188918 and MK188919, respectively. An alignment of WTK1 KinI and KinII domains was performed with ClustalW software (Thompson *et al.*, 1994). Secondary structures of kinase-like and pseudokinase-like domains were obtained using web server “PredictProtein” (Rost *et al.*, 2004). Key conserved residues, ATP binding site, catalytic loop and activation loop were defined as previously described (Klymiuk *et al.*, 2018).

## Results

### *YrH52* and *YrG303* express distinct phenotypes in response to *Pst* inoculation

*YrH52* in WEW accession H52 background provides a strong resistance response (IT1) to inoculation with the *Pst* isolate #5006 displaying only small dots or spots of HR in sites of fungal penetration and initial establishment of fungal colonies (**Fig. 1**). The HR area corresponds to the size of fungal colonies visualized by fluorescent microscopy with WGA-FITC fluorescent dye (**Fig. 1**). In general, the sizes of colonies as well as amount of HR needed to stop fungal development in *YrH52* were much smaller than these of *Yr15* in WEW backgrounds (**Fig. 1**).

*YrG303* in the WEW donor background in most cases (7 out of 10 tested plants) displayed moderate resistance response (IT5) to inoculation with *Pst* isolate #5006 characterized by the chlorotic areas, indicating an extensive hypersensitive response (HR), accompanied by some level of sporulation (**Fig. 1**). Such an intermediate level of resistance response of *YrG303* is distinct from the strong responses of *Yr15* and *YrH52* (**Fig. 1**). Furthermore, fluorescence microscopy revealed development of a massive net of fungal hyphae and some uredinia bags with spores in WEW G303 plants inoculated with *Pst* (**Fig. 1**). However, it should be noted that some G303 plants showed IT1, IT3 and IT4 (1 out of 10 tested plants for each mentioned ITs).

The tetraploid *YrG303* F_2_ mapping population showed the expected phenotypic segregation for a single dominant resistance gene 3:1 (χ^2^=1.47; *P*=0.1) in response to *Pst* inoculation based on phenotyping of 843 F_2_ plants, and F_1_ plants exhibit full resistance with IT1. We used the IT scale of 0 to 9 (Line and Qayoum, 1992), based on which G303 resistant parent showed IT1-5 and D447 susceptible parent showed IT8-9. Thus, phenotypic responses of F_2_ plants with IT1-6 were classified as resistant, while with IT7-9 were classified as susceptible. To confirm moderate resistance phenotypic responses (IT4-6), all homozygous recombinants were phenotyped in field trials at adult stage at F_4-5_ generation (**Fig. S1**).

The fine mapping of *YrG303* demonstrated that 3 out of the 124 homozygous recombinant lines were susceptible at the seedling stage even though they were expected to be resistant based on their genotype. (**Fig. S1**). We repeated seedling inoculation experiments in multiple generations (F_3_-F_5_) with similar results. However, artificial field inoculation with the same *Pst* isolate at the adult stage resulted in resistance response of these lines (**Fig. 2**). Furthermore, G303 plants exhibited different levels of resistance responses upon inoculation at the seedling stage (IT1-5) as compared to complete resistance (IT0-1) at the adult stage. In addition, after introgression of *YrG303* to the hexaploid common wheat cultivar Avocet (introgression line 2298), this gene provided full resistance at seedling stage to *Pst* inoculation with IT1 (**Fig. 3**).

**Figure 2.**
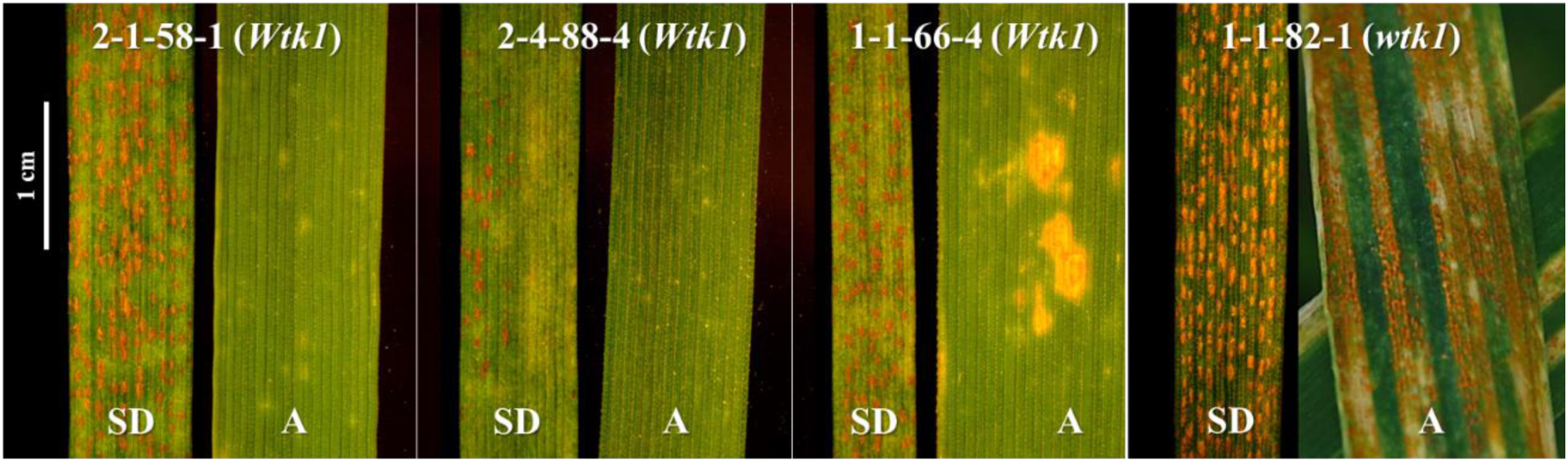
Comparisons of seedling and adult responses of four recombinant lines from the *YrG303* segregating mapping population (G303×D447) to inoculation with *Pst* isolate #5006 at 14 and 21 dpi, respectively. SD – seedling inoculation under controlled growth chamber conditions, A – adult inoculation under field conditions. RILs 2-1-58-1, 2-4-88-4, and 1-1-66-4 harbor the functional *Wtk1* allele, while 1-1-82-1 harbors the non-functional *wtk1* allele, and was used as a susceptible control. This figure is available in colour at JXB online.

**Figure 3.**
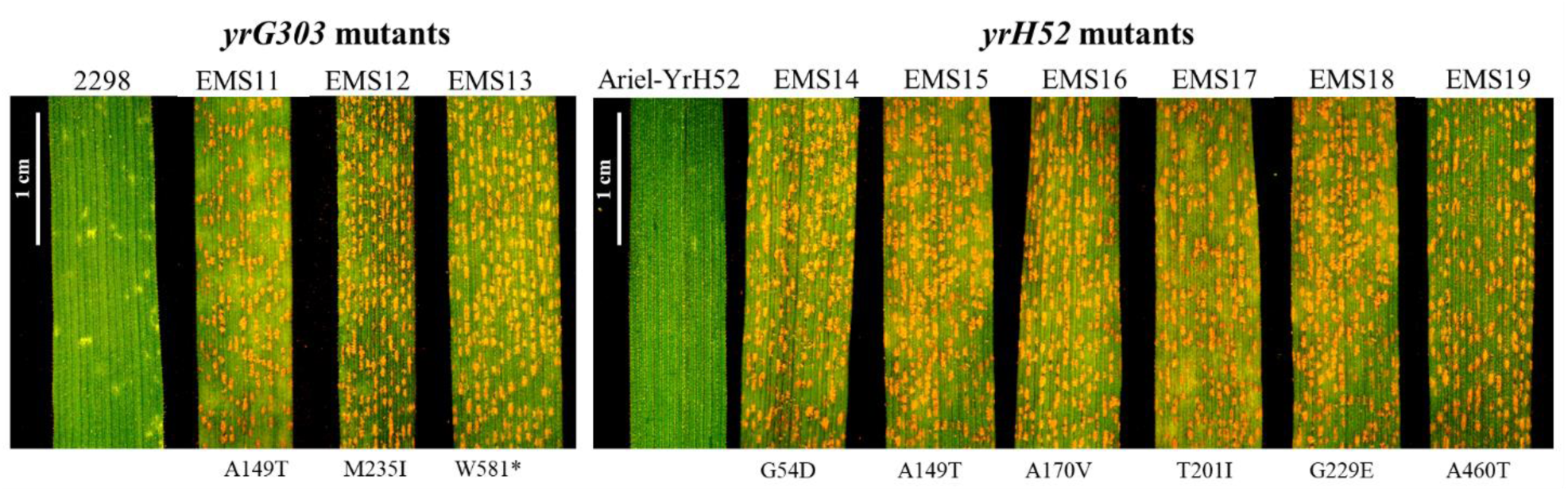
Susceptible reaction of *yrG303* and *yrH52* mutants to *Pst* inoculation at 14 dpi with *Pst* isolate #5006. 2298 and Ariel-YrH52 are wild type *YrG303* and *YrH52* hexaploid introgression lines, respectively, used to develop mutants. This figure is available in colour at JXB online.

### Fine mapping of *YrH52* and *YrG303*

#### YrH52

Previously, *YrH52* was mapped using low-resolution mapping as proximal to *Yr15* (Peng *et al.*, 2000a,b). A high-resolution genetic map of the *YrH52* genetic region was constructed using an F_2_ mapping population. In total, 3,549 F_2_ plants of the *YrH52* mapping population were screened for recombinants using the *YrH52* flanking markers, of which 3,211 F_2_ plants were screened using *wmc406* and *gwm413* markers (218 recombination events were detected) and 338 F_2_ plants were screened with *barc8* and *gwm273* markers (12 recombination events were detected). In total, 194 homozygous F_3-4_ RILs were developed and used for high-resolution mapping based on the graphical genotypes approach (**Fig. 4**). The genetic region between *barc8* and *gwm273* markers was saturated with ten CAPS and eight KASP markers (**Fig. S2**). Fine mapping of *YrH52* revealed that its genetic position co-segregated with three dominant (*uhw292, uhw300*, and *uhw301*) and three co-dominant markers (*uhw297, uhw296*, and *uhw259*) and overlaps with the location of the *Yr15* locus (Klymiuk *et al.*, 2018). Sequencing of the full-length genomic *WTK1* from WEW H52 revealed that it is identical to *Wtk1* from WEW G25.

**Figure 4.**
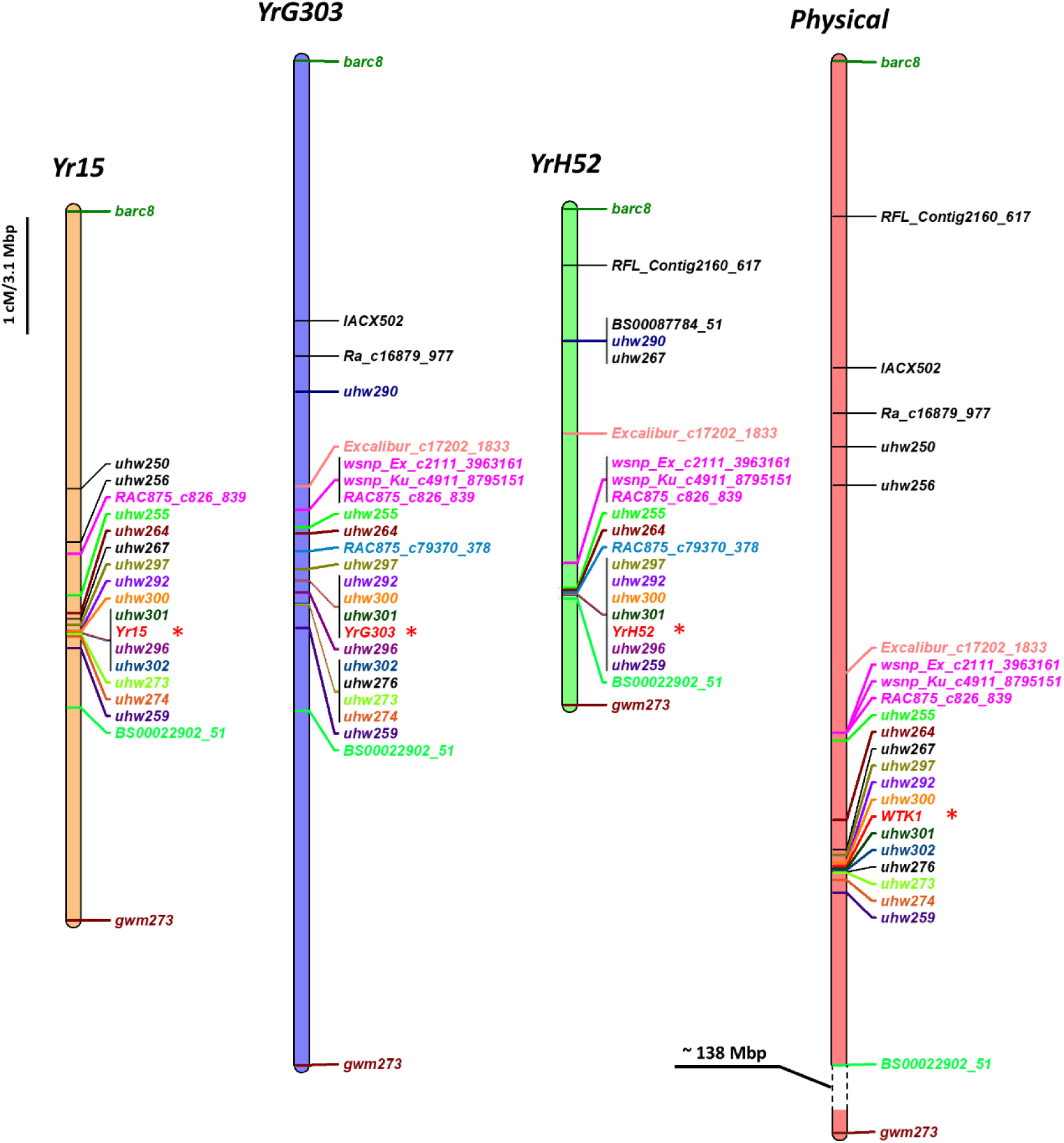
Genetic and physical maps of *Yr15, YrG303* and *YrH52* showing the same position for all three genes that correspond to *WTK1*. The consensus physical map represents three reference genomes, based on 1BS pseudomolecules of WEW Zavitan, *T. durum* Svevo, and *T. aestivum* CS. The consensus physical map contains only collinear markers between the genetic and the physical maps. This figure is available in colour at JXB online.

#### YrG303

We conducted preliminary experiments and localized *YrG303* distal to marker *gwm413*, suggesting that *YrG303* is different from *Yr15*, which was mapped proximal to *gwm413* (Yaniv *et al.*, 2015). In order to localize *YrG303* more precisely relative to *Yr15*, we performed fine mapping of this locus. A high-resolution genetic map was developed by screening of 1,381 F_2_ plants, from the *YrG303* tetraploid mapping population, with SSR markers *gwm273* and *barc8* flanking a chromosome interval of 8.5 cM, and then 536 F_2_ plants with internal SNP markers *RAC875_c826_839* and *BS00022902_51*, flanking a 1.7 cM interval that harbors the target gene (**Fig. 4**). These screenings identified heterozygous F_2_ recombinant lines within the region spanning *YrG303*. From them, a total of 124 homozygous F_3-4_ recombinants were developed and used for graphical genotyping of *YrG303*. These recombinants were genotyped with 23 PCR markers (SSR, CAPS, KASP, and dominant gene-specific markers) that showed polymorphisms between the parental lines (G303 and D447) (**Fig. 4; Fig. S1**) and phenotyped by *Pst* inoculation. Three dominant gene-specific markers (*uhw292, uhw300*, and *uhw301*), previously mapped to chromosome 1BS (Klymiuk *et al.*, 2018), were found to be co-segregating with the *YrG303 Pst* resistance phenotype and verified the location of *YrG303* on the short arm of chromosome 1B. The fine mapping of the gene revealed that the position of *YrG303* was the same as the *Yr15* locus (Klymiuk *et al.*, 2018) (**Fig. 4**). Sequencing of the full-length genomic *WTK1* from WEW G303 individual plant showing IT1, and another one showing IT5, revealed full identity between the two sequences and *Wtk1* from WEW G25.

### Validation of *WTK1* as a candidate gene for *YrG303* and *YrH52* resistance genes by EMS mutagenesis

A total of ∼2500 seeds of *YrG303* introgression lines and ∼1000 seeds of *YrH52* introgression line were treated with EMS. The germination rate of M_1_ seeds ranged between 73 to 80%. For *YrG303*, we screened approximately 800 M_1_-derived families generated from F_6_ RILs of the hexaploid A95 population, and only one susceptible mutant was identified. Furthermore, we screened approximately 1,200 M_1_-derived families generated from hexaploid line 2298, and two additional susceptible mutants were identified. Sequencing of *WTK1* from these susceptible lines revealed that they all carried independent mutations in the coding region of *WTK1* sequence (**Fig. 3, Table S3**). For *YrH52*, we screened 724 M_1_-derived families generated from the hexaploid introgression line Ariel-YrH52 and identified six mutants that all contained independent mutations in *WTK1* (**Fig. 3, Table S3**).

### Point mutations result in disruption of WTK1 function

WTK1 possesses two distinct kinase domains with the first kinase domain (KinI) predicted to encode a functional kinase, while the second (KinII) lacks several critical residues and therefore is predicted to be a pseudokinase (Klymiuk *et al.*, 2018). All 19 susceptible EMS mutants developed based on *Yr15* (Klymiuk *et al.*, 2018), *YrH52* and *YrG303* introgression lines were analyzed for the positions of specific mutations within *Wtk1* (**Fig. 5**). Five susceptible *yr15* mutants carried SNP changes in KinI kinase domain and five carried SNP changes in KinII pseudokinase domain (Klymiuk *et al.*, 2018). The majority of the non-functional *yrG303* and *yrH52* mutants developed within the current study carried mutations in the KinI; only one *yrG303* mutant and one *yrH52* mutant carried mutations in KinII pseudokinase domain (**Table S3**). Resistant sister lines of all *yrG303* mutants deriving from the same M_1_ families showed the presence of the wild type allele of *Wtk1*, therefore validating that the mutations in KinI and KinII domains are responsible for the loss-of-function, and that the resistance conferred by *Yr15, YrG303* and *YrH52* is encoded by only one gene.

**Figure 5.**
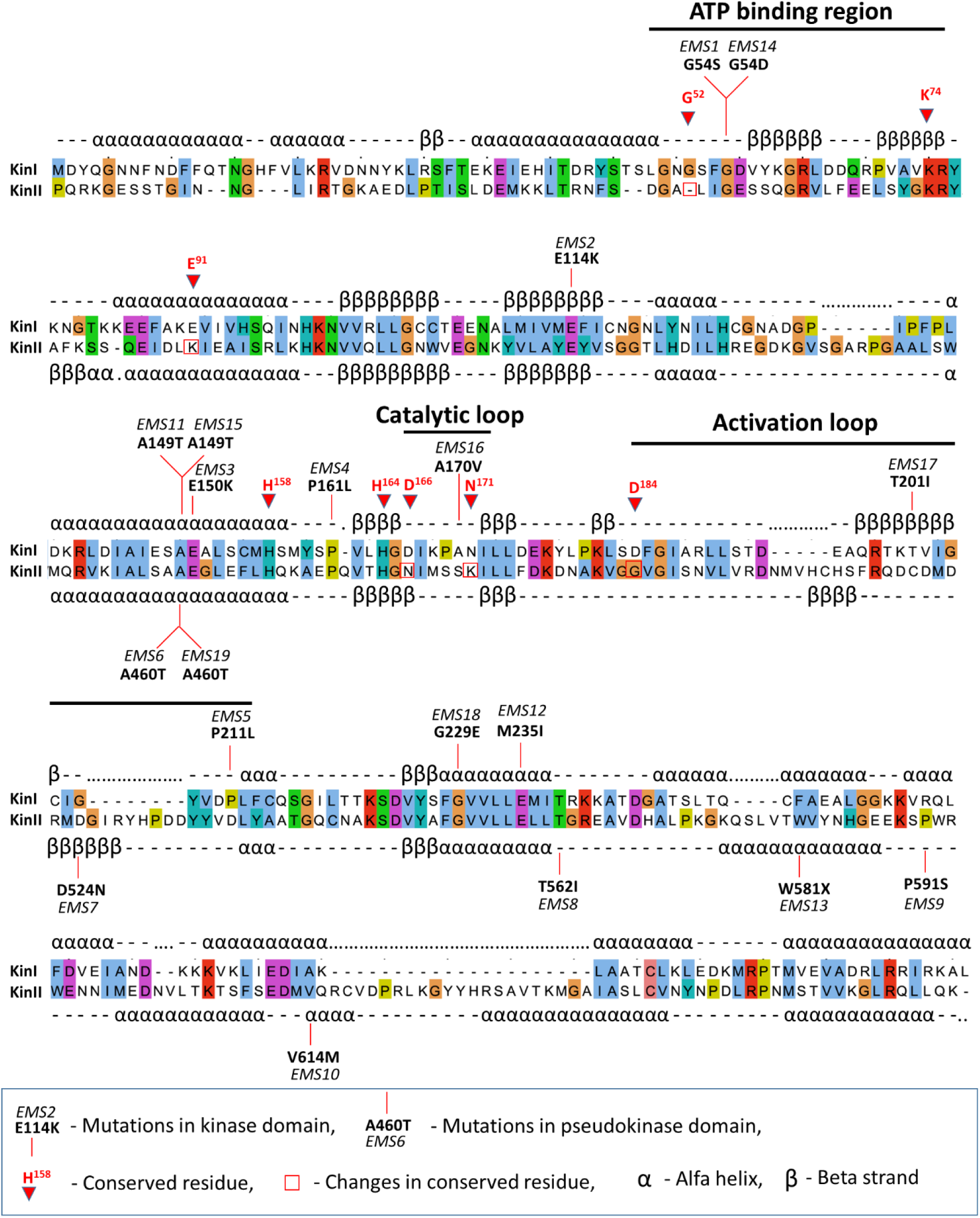
Primary and secondary structures of WTK1 kinase and pseudokinase domains alongside with positions of knock-out EMS mutations in *yr15, yrG303* and *yrH52* susceptible mutants. The diagram of WTK1 domain architecture highlights eight key conserved residues in the kinase domain (with numbers that correspond to their positions in cAPK (Hanks *et al.*, 1988)) and the absence of five of them in the pseudokinase domain. Red lines indicate EMS mutations that block resistance. KinI = kinase domain, Kin II = pseudokinase domain. This figure is available in colour at JXB online.

The EMS mutations in WTK1 that affect recognition in *yr15, yrG303* and *yrH52* susceptible mutants have not been identified in key conserved residues, but do map to conserved regions in critical kinase subdomains such as the ATP binding region, catalytic loop and activation loop (**Fig. 5**). Three amino acids (G^54^, A^149^ and A^460^) were disrupted in different wheat genetic backgrounds and all inhibited WTK1-mediated immunity to yellow rust, confirming that these residues are critical for WTK1 responses. (**Fig. 5**). G^54^ is located in KinI’s ATP binding region and may directly affect WTK1’s kinase activity (**Fig. 5**). A^149^ and A^460^ map to the same alpha-helical region present in KinI and KinII domains. Neither A^149^ nor A^460^ map to catalytic residues, but their presence in the same region of KinI and KinII indicates that these alanine residues may be required for proper WTK1 folding and/or *Pst* effector binding.

## Discussion

Recent advances in plant innate immunity system have shed light on possible mechanisms and organization of resistance networks. However, identification of R-genes and an understanding of their allelic series serve as an essential initial step to dissect complex plant-pathogen interactions. Many studies have been performed to discover novel *Pst* R-genes. However, the deployment of novel R-genes from WEW into common wheat is a long process due to ploidy differences, negative linkage drag, etc. For R-genes that map to similar chromosome regions, it is important to determine if they are in fact different genes or alleles of the same genes because this will impact strategies for their effective deployment.

### *YrG303, YrH52* and *Yr15* R-genes derived from WEW

*Yr15, YrG303* and *YrH52* genes originated from different WEW accessions and were all mapped to the short arm of chromosome 1B. However, they were initially considered distinct R-genes, due to differences in geographic distribution of their donor lines and different resistance reaction patterns in response to inoculation with the same *Pst* races. *Yr15* originated in WEW accession G25 collected in Rosh Pinna, Israel (Gerechter-Amitai *et al.*, 1989), while WEW accession G303 harboring *YrG303* was collected in Dishon (Israel), and WEW accession H52, the donor line of *YrH52*, was collected in Mt. Hermon (Israel), that represent different habitats. These three WEW accessions showed different resistance profiles, such as susceptibility of G25 to *Pst* isolate #81065 from Afghanistan and resistance of G303 to the same isolate (Van Silfhout, 1989); as well as different level of resistance to *Pst* isolate #5006 from Israel with *YrH52* (H52) displaying IT1, *Yr15* (G25) IT3 and *YrG303* (G303) IT5, all proved to have corresponding differences at the level of development of pathogenic fungal colonies (Fig. 1). Fine genetic mapping of the three genes demonstrated that all three map to the same genetic interval on chromosome arm 1BS (Fig. 4, Fig. S1, Fig. S2). Moreover, sequencing of *WTK1* from the susceptible *yrG303* and *yrH52* EMS mutants showed that all of them contain loss-of-function mutations in *Wtk1* (Fig. 3, Fig. 5, Table S3). Thus, other genes reported to be located on 1BS chromosome, especially these originating from WEW, such as *YrSM139-1B* (Zhang *et al.*, 2016) and *YrTz2* (Wang *et al.*, 2018), should be tested for allelism/identity to *Wtk1*.

The sequences of *Wtk1* from G25, G303 and H52 have been published previously (Klymiuk *et al.*, 2018; Klymiuk *et al.*, 2019b), however in G303 and H52 genetic backgrounds we did not prove that the resistance phenotype is conferred by the *Wtk1* allele. Here, based on fine genetic mapping, we show that the genetic positions of *YrG303* and *YrH52* indeed coincide with the physical position of *Wtk1* (Fig. 4), and that mutations in *Wtk1* lead to susceptibility in both *YrG303* and *YrH52* introgression lines (Fig. 3, Fig. 5, Table S3). Furthermore, *YrG303* and *YrH52* both possess a *Wtk1* functional allele that is identical to the *Yr15* sequence of *Wtk1*. We hypothesize that such high sequence conservation represents a specific character of TKP proteins as compared with the high level of variability observed for the known nucleotide-binding domain and leucine-rich repeat (NLR) proteins (MacQueen *et al.*, 2019). Clusterization, rapid evolution and the presence of multiple alleles is a known character of many NLRs (Bhullar *et al.*, 2009; Marchal *et al.*, 2018; Adachi *et al.*, 2019). TKPs also tend to cluster together, probably reflecting the evolutionary mechanisms by which they evolved via duplication or fusion of two kinase domains (Klymiuk *et al.*, 2018). For example, WTK1 has three tandem copies on each chromosomes 6A and 6B (Klymiuk *et al.*, 2019b). However, in contrast to NLRs, we did not detect diverse *Wtk1* alleles, and differences in phenotypic responses are likely associated to differences in the genetic backgrounds as discussed below.

### The spectrum of *Wtk1* phenotypic responses

Three components, host, pathogen and environment, serve as a base for the concept of disease triangle and determine the degree of severity of the disease (Agrios, 2005). According to the gene-for-gene model, host-pathogen interactions result in co-evolution of virulence genes (e.g. pathogen effectors) and resistance genes (e.g. host receptors) (Flor, 1971). However, even in case of compatible interaction between host and pathogen, disease may not occur or symptoms will be limited due to unfavorable environmental conditions for pathogen development (Agrios, 2005). Plant immunity system is multifaceted and comprises recognition (via receptors) and response (a transduction network of multiple genes deployed) parts (Jones & Dangl, 2006). Different alleles of various R-genes, e.g. wheat *Yr5*/*YrSP* (Marchal *et al.*, 2018), wheat *Pm3* (Srichumpa *et al.*, 2005), barley Mla (Seeholzer *et al.*, 2010), flax L locus (Ellis *et al.*, 1999), tomato *Pto* (Rose *et al.*, 2005), Arabidopsis *Rpm1* locus (Stahl *et al.*, 1999), Arabidopsis *RPP13* (Rose *et al.*, 2004), possess distinct resistance specificities and responses. These are based on the possibility of allelic variation in R-genes to recognize pathogen effectors with different efficiencies. Clearly, this does not apply for *Yr15, YrG303* and *YrH52* because their sequences are identical.

One possible explanation of observed variation in phenotypic responses of *Wtk1*-carriers could be related to differences in expression levels. However, we have shown previously (Klymiuk *et al.*, 2018) that differences in expression levels of *Yr15* (*Wtk1*) in transgenic and introgression lines did not relate to phenotypic expression differences. Also the basal expression of *Yr15* (*Wtk1*) expression is low and did not change significantly after inoculation with *Pst* (Klymiuk *et al.*, 2018).

Introgression of these genes into different genetic backgrounds shows repeatable differences in resistance response upon *Pst* inoculation. We have previously shown that tetraploid (*T. turgidum* ssp. *durum*) and hexaploid (*T. aestivum*) *Yr15* introgression lines all exhibit diverse phenotypic responses to *Pst* inoculation (Klymiuk *et al.*, 2018). We demonstrated that the introgression of *YrG303* from WEW G303 to *T. aestivum* 2298 improved the resistance phenotype from IT5 to IT1 (Fig. 1, Fig. 3). This could be possible due to variable regulation of *Wtk1* expression, perhaps because of variation in the flanking regulatory sequences (which were not obtained in the current study). However, this seems unlikely given that the introgression lines mentioned here were developed through conventional breeding and likely carry relatively large introgressed segments, including those flanking *Wtk1* up- and down-streams (Yaniv *et al.*, 2015). Similar situations occur during introgression or transformation of other R-genes. For example, transformation of *LR34res* allele into two genetic backgrounds of wheat resulted in variation in resistance of transgenic lines, which was explained by differences in genes modifying *LR34res* activity (Risk *et al.*, 2012). Moreover, diverse phenotypic responses are typical not only for wheat-rusts interactions, but also for other pathosystems. In particular, transfer of R-genes/quantitative resistance loci for resistance to rice blast resulted in variation in resistance reaction of improved lines to inoculation with *Magnaporthe oryzae* (Hasan *et al.*, 2016). A more likely explanation of unexpected phenotypic results is that effectiveness of R-genes is altered following introgression into a new background (Adachi *et al.*, 2019). According to current understanding of plant immunity system, some response networks may be shared between different receptors, while others are very specific. Faulty connections between nodes in the NLR network may arise after a cross between distinct plant genotypes that can result in NLR mis-regulation and autoimmunity of progeny, although in parental lines the immune system worked well (Adachi *et al.*, 2019).

Although *Wtk1* provides resistance at both, seedling and adult stages, *YrG303* exhibited a resistance response in three independent tetraploid recombinant lines only at the adult stage under field inoculation (Fig. 2). Taking into account that controlled conditions in dew chamber used for seeding inoculation are more favorable for *Pst* development than field conditions used for adult stage inoculation, our results suggest that the environment plays an important role in the resistance response of these genes. Moreover, WEW accessions that carry functional *Wtk1* alleles and were subjected to natural selection and adaptation (Klymiuk *et al.*, 2019b), show different levels of resistance within the same experiment. An analogous situation was shown for *Pto* locus in wild tomato (*Lycopersicon* spp.) populations, where susceptible phenotypes were detected even under presence of alleles conferring *AvrPto* recognition, suggesting non-functionality of some of the genes from *Pto* resistance response network (Rose *et al.*, 2005). Taking all of the above into account, we hypothesize that phenotypic differences in the presence of *Wtk1* likely originate from differences in the genetic background, rather than from the presence of different *Wtk1* resistant alleles. Nevertheless, we do not exclude that other factors, e.g. presence of genes-suppressors of R-genes in some genetic backgrounds, may influence phenotypic responses of *Wtk1*-carrier lines (Kema *et al.*, 1995).

### Positions of loss-of-function mutations in WTK1

Both the kinase and pseudokinase domains of WTK1 are necessary to provide resistance to *Pst* (Klymiuk *et al.*, 2018). Plant pseudokinases are important players in diverse biological processes and represent ∼10% of the kinase domains in higher eukaryotes (Castells and Casacuberta, 2007; Reiterer *et al.*, 2014; Niu *et al.*, 2016). Emerging evidence indicates that pseudokinases themselves act as signaling molecules and modulate the activity of catalytically active kinase partners (Müller *et al.*, 2008; Reiterer *et al.*, 2014). Consistent with this hypothesis and the previous study (Klymiuk *et al.*, 2018), EMS mutations blocking resistance were mapped to both pseudokinase and kinase domains of WTK1 (Fig. 5). In the current study, we identified informative EMS mutations in conserved kinase motifs such as the ATP binding region, the catalytic loop and the activation loop (Kornev and Taylor, 2010). Nevertheless, these mutations did not affect previously defined key conserved residues involved in kinase activity (Kannan *et al.*, 2007; Klymiuk *etal.*, 2018). It seems that these mutations are still able to affect WTK1’s kinase activity possibly by disrupting the correct folding of the protein or by preventing binding of pathogen effectors. Interestingly, we identified single nucleotide mutations caused by EMS treatment in three residues, G^54^, A^149^ and A^460^, that each blocked resistance in different genetic backgrounds (Fig. 5). G^54^ maps to KinI’s ATP binding region and corresponds to glycine G^55^ of the α cAMP-dependent protein kinase catalytic subunit (cAPK) (Hanks *et al.*, 1988) known to be part of a Glycine-rich Loop that coordinates the ATP phosphates during binding (Taylor and Kornev, 2011). A^149^ and A^460^ occur in the same alpha-helical region in both KinI and KinII domains of WTK1, respectively, and correspond to A^151^ of cAPK (Hanks *et al.*, 1988). Although the functions of A^151^ itself remain unclear, it is located close to LXXLH^158^ motif that is involved in providing the link between the catalytically important DFG motif and substrate binding regions as a part of hydrogen-bonding network (Kannan *et al.*, 2007). Thus, it seems that the A^149^/A^460^ region of WTK1 is important for KinI/II associations, effector binding or target binding.

### Conclusions and future perspectives

Here we show that *Yr15, YrG303* and *YrH52* all represent one tandem kinase gene, *Wtk1. Wtk1* sequences of *YrG303* and *YrH52* are identical to those of *Yr15*. Differences in phenotypic responses of WEW donor lines and introgression lines of *Yr15, YrG303* and *YrH52* are probably related to differences due to genetic backgrounds rather than to the presence of different alleles. Our results indicate that *YrG303* and *YrH52* are synonymous to *Yr15*. However, we cannot exclude the option that the three WEW donor lines (G25, H52 and G303) harbor additional *Yr* genes. Positions of mutations in EMS treated lines that led to disruption of *Wtk1* function were associated with conserved regions in critical kinase subdomains, although key conserved residues were not affected. This information will be useful for future work on the possible molecular mechanism of *Wtk1* and its role in plant innate immunity. The *Wtk1*-mediated resistance network is diverse in WEW natural populations subjected to natural selection and adaptation, confirming that WEW natural populations have potential to serve as a good source for evolutionary studies of different traits and multifaceted gene networks.

## Abbreviations

cAPK: α cAMP dependent protein kinase catalytic subunit
CAPS: cleaved amplified polymorphic sequences
dpi: days post inoculation
EMS: ethyl methanesulfonate
HR: hypersensitive response
KASP: kompetitive allele specific PCR
NLR: nucleotide-binding domain and leucine-rich repeat
*Pst*: *Puccinia striiformis* f. sp. *tritici*
R-genes: resistance genes
SNP: single nucleotide polymorphism
SSR: simple sequence repeat
TKP: Tandem Kinase-Pseudokinase
WEW: wild emmer wheat
WGA: wheat germ agglutinin
*wtk1*: and non-functional *WTK1* allele
*Wtk1*: functional *WTK1* allele
WTK1: Wheat Tandem Kinase 1
*Yr* gene: yellow rust resistance gene

## Supplementary data

**Table S1.** A list of SSR, CAPS and KASP markers used in the current study.

**Table S2.** A list of KASP markers from the *Yr15* region developed based on SNPs from the wheat 15K SNP array.

**Table S3.** Molecular characterization of the *yr15, yrG303* and *yrH52* EMS mutants.

**Fig. S1.** Graphical genotype of selected recombinant lines (RILs) from *YrG303* tetraploid mapping population.

**Fig. S2.** Graphical genotype of selected RILs from *YrH52* mapping population.

## Acknowledgements

This research was supported by the US-Israel Binational Agricultural Research and Development Fund (IS-4628-13, US-4916-16), and the Israel Science Foundation (1719/08, 1366/18). We thank S. Barinova, J. Cheng, T. Kis-Papo, S. Khalifa, T. Krugman, D. Lewinsohn, I. Manov, O. Rybak, I. Shams for their support, advice and discussions.

