## Supplementary Information for "Three previously characterized resistances to yellow rust are encoded by a single locus *Wtk1*"

**Table S1.** A list of SSR, CAPS and KASP markers used in the current study.

| Name of marker | Type of marker | Primer sequence | Reference |
|---|---|---|---|
| *gwm413* | SSR | TGCTTGTCTAGATTGCTTGGG<br>GATCGTCTCGTCCTTGGCA | Röder *et al.* (1998) |
| *wmc406* | SSR | TATGAGGGTCGGATCAATACAA<br>CGAGTTTACTGCAAACAAATGG | Somers *et al.* (2004) |
| *barc8* | SSR | GCGGGAATCATGCATAGGAAAACAGAA<br>GCGGGGGCGAAACATACACATAAAAACA | Song *et al.* (2005) |
| *uhw290** | CAPS | ATAGCAGGATCGCAGCAAAA<br>GAGGTATAACAATGTGATCGTTCTG | Current study |
| *uhw250* | EST | CTGCTCACTTTTTGCCTGTG<br>AAAAGTTGTTGCTCTGCTTTT | Klymiuk *et al.* 2018 |
| *uhw256* | EST | GTTACCCTCCACAGCAAGGT<br>GCGCATTACTTCCACTTCTTG | Klymiuk *et al.* 2018 |
| *RAC875_c826_839* | KASP | ACGAAGGTTCTGTTTTCACCA<br>ACGAAGGTTCTGTTTTCACCG<br>TCTTCTTGCTCAAAGGTAAGAGT | Klymiuk *et al.* 2018 |
| *uhw255* | CAPS | GATGCTCTGCACATGTGTTATG<br>GCAGCTCCAGCTTATTCGTC | Klymiuk *et al.* 2018 |
| *uhw264* | CAPS | GGTCTCTTGCAACATACAGTAACAA<br>GAGTGGTAGTCTAGTAGAGGTTGGTG | Klymiuk *et al.* 2018 |
| *uhw267* | CAPS | TGGTAATCAAGTTTCACATTGTTCA<br>GGAAGGACACCTTTCGGTATT | Klymiuk *et al.* 2018 |
| *uhw297* | CAPS | CAGATGACCAACCAAAAGCA<br>GTCATATTGGTGCCCAGTGA | Klymiuk *et al.* 2018 |
| *uhw292* | dominant | GACTTTCTTCCCTCGGGACT<br>CTCGCACGCCTATAAAAGGA | Klymiuk *et al.* 2018 |
| *uhw300* | dominant | CCGTGTCAGCCACCTACAAT<br>GCACTCTACCACCGAACACA | Klymiuk *et al.* 2018 |
| *uhw301* | dominant | GTAGTGGCTCGTTCGGTGAT<br>TTTCGCATCCCACCCTACTG | Klymiuk *et al.* 2018 |
| *uhw296* | CAPS | CAACCGTGCCTCCAAACA<br>CGGGTGTTGTCCGTTGAG | Klymiuk *et al.* 2018 |
| *uhw302* | dominant | CATCCATTCCTCCGACAAGT<br>CACTGCAATGCAAAAATGCT | Klymiuk *et al.* 2018 |
| *uhw276* | CAPS | TCTGTGATGCCTGTGATGGT<br>AAAGTTTGGGATTTGGCAAT | Klymiuk *et al.* 2018 |
| *uhw273* | CAPS | GGTGACGGCGAGTGTACG<br>GACGCAATTGTCCGCTGT | Klymiuk *et al.* 2018 |
| *uhw274* | CAPS | AAGCTCCGCTGCAATGAC<br>ACCTGACATCCTCGAACCAC | Klymiuk *et al.* 2018 |
| *uhw259* | CAPS | CTGTATTCTAATGCAGATTAGCTGTT<br>CACGCATAATTTTGTCCACAC | Klymiuk *et al.* 2018 |
| *BS00022902_51* | KASP | ATGTGCGGCAGGAGAAGA<br>ATGTGCGGCAGGAGAAGG<br>ATACTCTTCACGGTCGTCTTC | Klymiuk *et al.* 2018 |
| *gwm273* | SSR | ATTGGACGGACAGATGCTTT<br>AGCAGTGAGGAAGGGGATC | Röder *et al.* (1998) |

[*] Temperature of elongation 59°C; Enzyme *Aci*I; Fragment sizes: *T. dicoccoides* H52 = 266 bp, 148 bp; *T. durum* cv. Langdon = 414 bp.

**Table S2.** A list of KASP markers from the *Yr15* region developed based on SNPs from the wheat 15K SNP array.

| Name of marker | Primer A | Primer B | Primer Common |
|---|---|---|---|
| *RFL_Contig2160_617* | cgtgtgaatgtgtactctacca | cgtgtgaatgtgtactctaccg | ggacccagaatgcccagtg |
| *IACX502* | cacgcctaacaaaacatatccatt | cacgcctaacaaaacatatccatc | tgggagttcttatttaatctttcgg |
| *Ra_c16879_977* | cagagcaaaagccatcaatctta | cagagcaaaagccatcaatcttc | tctgagaaagatgccagaacg |
| *BS00087784_51* | gagctgacagatgggggt | gagctgacagatgggggc | ccatgacatcagaaaagttgatgat |
| *Excalibur_c17202_1833* | gttagagcccaattttcaagcta | gttagagcccaattttcaagctg | aggagcttatattgctatgtacagt |
| *wsnp_Ku_c4911_8795151* | gttcatcaactgttgagctgtt | gttcatcaactgttgagctgtc | gctgctgtagtttctgattgtg |
| *wsnp_Ex_c2111_3963161* | agcagcatacgattaactcagtt | agcagcatacgattaactcagtc | ctgttggcgcagaagctg |
| *RAC875_c79370_378* | tggaactgatggtgtccagt | tggaactgatggtgtccagc | gcggcatcaacttccccg |

**Table S3.** Molecular characterization of the *yr15*#, *yrG303* and *yrH52* EMS mutants

| Domain | Base substitution* | Effect on amino acid† | Gene | Mutant No. | Name of mutant |
|---|---|---|---|---|---|
| KinI | G 160 A | G 54 S | *yr15* | EMS1 | Suncea+*yr15*-L18 |
| KinI | G 161 A | G 54 D | *yrH52* | EMS14 | M52-4 |
| KinI | G 340 A | E 114 K | *yr15* | EMS2 | Suncea+*yr15*-L89 |
| KinI | G 445 A | A 149 T | *yrG303* | EMS11 | M2298-410-16 |
| KinI | G 445 A | A 149 T | *yrH52* | EMS15 | M52-9 |
| KinI | G 448 A | E 150 K | *yr15* | EMS3 | B9+*yr15*-L1351 |
| KinI | C 482 T | P 161 L | *yr15* | EMS4 | Avocet+*yr15*-1 |
| KinI | C 509 T | A 170 V | *yrH52* | EMS16 | M52-14 |
| KinI | C 602 T | T 201 I | *yrH52* | EMS17 | M52-18 |
| KinI | C 632 T | P 211 L | *yr15* | EMS5 | Avocet+*yr15*-L90 |
| KinI | G 686 A | G 229 E | *yrH52* | EMS18 | M52-2 |
| KinI | G 705 A | M 235 I | *yrG303* | EMS12 | A95-126 |
| KinII | G 2922 A | A 460 T | *yr15* | EMS6 | Avocet+*yr15*-13 |
| KinII | G 2922 A | A 460 T | *yrH52* | EMS19 | M52-8 |
| KinII | G 3114 A | D 524 N | *yr15* | EMS7 | Excalibur+*yr15*-6L306 |
| KinII | C 3229 T | T 562 I | *yr15* | EMS8 | Avocet+*yr15*-L72 |
| KinII | G 3281 A | W 581 * | *yrG303* | EMS13 | M2298-767-16 |
| KinII | C 3315 T | P 591 S | *yr15* | EMS9 | B9+*yr15*-LF |
| KinII | G 3469 A | V 614 M | *yr15* | EMS10 | Excalibur+*yr15*-L137 |

#Full description of *yr15* mutants is provided in Klymiuk *et al.* (2018).

*The first letter indicates the wild-type nucleotide, the number indicates its position relative to the ATG start codon, and the last letter shows the mutant nucleotide. The complete WTK1 coding regions of the above 19 mutants were sequenced; no additional mutations were detected.

†The first letter indicates the wild-type amino acid, the number indicates its position relative to the start methionine, and the last letter shows the mutant amino acid.

| #RIL | *barc8* | *IACX502* | *Ra_c16879_977* | *uhw290* | *Excalibur_c17202_1833* | *wsnp_Ku_c4911_8795151* | *wsnp_Ex_c2111_3963161* | *RAC875_c826_839* | *uhw255* | *uhw264* | *RAC875_c79370_378* | *uhw297* | *uhw292* | *uhw300* | *YrG303* seedling | *YrG303* adult | *uhw301* | *uhw296* | *uhw302* | *uhw276* | *uhw273* | *uhw274* | *uhw259* | *BS00022902_51* | *gwm273* |
|---|---|---|---|---|---|---|---|---|---|---|---|---|---|---|---|---|---|---|---|---|---|---|---|---|---|
| 2_6_38_3 | B | B | B | B | B | B | B | B | B | B | B | B | B | B | B | B | B | B | B | B | B | B | B | B | A |
| 1_1_82_1 | B | B | B | B | B | B | B | B | B | B | B | B | B | B | B | B | B | B | B | B | B | B | B | B | A |
| 1_1_84_5 | B | B | B | B | B | B | B | B | B | B | B | B | B | B | B | B | B | B | B | B | B | B | B | A | A |
| 2_2_92_10 | B | B | B | B | B | B | B | B | B | B | B | B | B | B | B | B | B | B | B | B | B | B | B | A | A |
| 2_6_50_1 | B | B | B | B | B | B | B | B | B | B | B | B | B | B | B | B | B | B | B | B | B | B | A | A | A |
| 2_3_51_3 | B | B | B | B | B | B | B | B | B | B | B | B | B | B | B | B | B | B | A | A | A | A | A | A | A |
| 2_4_25_3 | B | B | B | B | B | B | B | B | B | B | B | B | B | B | B | B | B | A | A | A | A | A | A | A | A |
| 1_3_19_4 | B | B | B | B | B | B | B | B | B | B | B | A | A | A | A | A | A | A | A | A | A | A | A | A | A |
| 2_1_12_10 | B | B | B | B | B | B | B | B | B | B | A | A | A | A | A | A | A | A | A | A | A | A | A | A | A |
| 1_2_66_10 | B | B | B | B | B | B | B | B | B | B | A | A | A | A | A | A | A | A | A | A | A | A | A | A | A |
| 2_4_23_5 | B | B | B | B | B | B | B | B | A | A | A | A | A | A | A | A | A | A | A | A | A | A | A | A | A |
| 2_1_54_2 | B | B | B | B | B | A | A | A | A | A | A | A | A | A | A | A | A | A | A | A | A | A | A | A | A |
| 2_3_32_4 | B | B | B | B | A | A | A | A | A | A | A | A | A | A | A | A | A | A | A | A | A | A | A | A | A |
| 1_2_70_1 | B | B | B | B | A | A | A | A | A | A | A | A | A | A | A | A | A | A | A | A | A | A | A | A | A |
| 3_2_74_4 | B | B | B | A | A | A | A | A | A | A | A | A | A | A | A | A | A | A | A | A | A | A | A | A | A |
| 2_1_64_1 | B | B | A | A | A | A | A | A | A | A | A | A | A | A | A | A | A | A | A | A | A | A | A | A | A |
| 2_4_88_4 | B | A | A | A | A | A | A | A | A | A | A | A | A | A | B | A | A | A | A | A | A | A | A | A | A |
| 3_2_64_2 | B | A | A | A | A | A | A | A | A | A | A | A | A | A | A | A | A | A | A | A | A | A | A | A | A |
| 2_2_60_2 | B | A | A | A | A | A | A | A | A | A | A | A | A | A | A | A | A | A | A | A | A | A | A | A | A |
| 1_1_66_4 | A | A | A | A | A | A | A | A | A | A | A | A | A | A | B | A | A | A | A | A | A | A | A | A | B |
| 2_7_4_3 | A | A | A | A | A | A | A | A | A | A | A | A | A | A | A | A | A | A | A | A | A | A | A | A | B |
| 3_3_48_3 | A | A | A | A | A | A | A | A | A | A | A | A | A | A | A | A | A | A | A | A | A | A | A | A | B |
| 2_1_58_1 | A | A | A | A | A | A | A | A | A | A | A | A | A | A | B | A | A | A | A | A | A | A | A | B | B |
| 2_1_59_2 | A | A | A | A | A | A | A | A | A | A | A | A | A | A | A | A | A | A | A | A | A | A | A | B | B |
| 1_1_79_1 | A | A | A | A | A | A | A | A | A | A | A | A | A | A | A | A | A | A | A | A | A | A | B | B | B |
| 2_5_74_8 | A | A | A | A | A | A | A | A | A | A | A | A | A | A | A | A | A | A | A | A | A | A | B | B | B |
| 2_5_80_4 | A | A | A | A | A | A | A | A | A | A | A | A | B | B | B | B | B | B | B | B | B | B | B | B | B |
| 1_3_58_9 | A | A | A | A | A | A | A | A | A | A | B | B | B | B | B | B | B | B | B | B | B | B | B | B | B |
| 2_4_45_3 | A | A | A | A | A | A | A | A | A | B | B | B | B | B | B | B | B | B | B | B | B | B | B | B | B |
| 3_2_5_3 | A | A | A | A | A | B | B | B | B | B | B | B | B | B | B | B | B | B | B | B | B | B | B | B | B |
| 2_3_30_2 | A | A | A | A | A | B | B | B | B | B | B | B | B | B | B | B | B | B | B | B | B | B | B | B | B |
| 3_3_10_2 | A | A | A | A | B | B | B | B | B | B | B | B | B | B | B | B | B | B | B | B | B | B | B | B | B |
| 3_2_26_2 | A | A | A | A | B | B | B | B | B | B | B | B | B | B | B | B | B | B | B | B | B | B | B | B | B |
| 3_2_86_2 | A | A | A | B | B | B | B | B | B | B | B | B | B | B | B | B | B | B | B | B | B | B | B | B | B |
| 3_2_18_3 | A | B | B | B | B | B | B | B | B | B | B | B | B | B | B | B | B | B | B | B | B | B | B | B | B |
| 2_2_7_4 | A | B | B | B | B | B | B | B | B | B | B | B | B | B | B | B | B | B | B | B | B | B | B | B | B |

**Fig. S1.** Graphical genotype of selected recombinant lines (RILs) from *YrG303* tetraploid mapping population. Allele A – resistant allele from *T. dicoccoides* acc. G303; allele B – susceptible allele from *T. durum* acc. D447. *YrG303* seedling and *YrG303* adult columns correspond to the phenotypes observed at seedling and adult screens with *Pst* (14 and 21 dpi, respectively). RILs marked in red showed different seedling and adult resistance response.

| #RIL | *barc8* | *RFL_Contig2160_617* | *BS00087784_51* | *uhw267* | *uhw290* | *Excalibur_c17202_1833* | *wsnp_Ku_c4911_8795151* | *RAC875_c826_839* | *wsnp_Ex_c2111_3963161* | *uhw255* | *uhw264* | *RAC875_c79370_378* | *uhw297* | *uhw292* | *uhw300* | *YrH52* | *uhw301* | *uhw296* | *uhw259* | *BS00022902_51* | *gwm273* |
|---|---|---|---|---|---|---|---|---|---|---|---|---|---|---|---|---|---|---|---|---|---|
| **V1_31_1** | B | B | B | B | B | B | B | B | B | B | B | B | B | B | B | B | B | B | B | B | A |
| **V1_60_6** | B | B | B | B | B | B | B | B | B | B | B | B | B | B | B | B | B | B | B | B | A |
| **GA19_37_12_1** | B | B | B | B | B | B | B | B | B | B | B | B | B | B | B | B | B | B | B | A | A |
| **GA_10_61_14_2** | B | B | B | B | B | B | B | B | B | B | B | B | A | A | A | A | A | A | A | A | A |
| **D11_86_6_4** | B | B | B | B | B | B | B | B | B | B | A | A | A | A | A | A | A | A | A | A | A |
| **GA_22_37_10_3** | B | B | B | B | B | B | B | B | B | A | A | A | A | A | A | A | A | A | A | A | A |
| **D16_5_11_2** | B | B | B | B | B | B | B | B | B | A | A | A | A | A | A | A | A | A | A | A | A |
| **V4_5_3** | B | B | B | B | B | B | B | B | B | A | A | A | A | A | A | A | A | A | A | A | A |
| **V2_90_4** | B | B | B | B | B | B | A | A | A | A | A | A | A | A | A | A | A | A | A | A | A |
| **D130_4_14_4** | B | B | B | B | B | B | A | A | A | A | A | A | A | A | A | A | A | A | A | A | A |
| **402_9_20_1_2** | B | B | B | B | B | B | A | A | A | A | A | A | A | A | A | A | A | A | A | A | A |
| **D128_47_7_4** | B | B | B | B | B | A | A | A | A | A | A | A | A | A | A | A | A | A | A | A | A |
| **V3_101_2** | B | B | B | B | B | A | A | A | A | A | A | A | A | A | A | A | A | A | A | A | A |
| **GA_26_27_6_2** | B | B | B | B | B | A | A | A | A | A | A | A | A | A | A | A | A | A | A | A | A |
| **d131_46_9_1** | B | B | A | A | A | A | A | A | A | A | A | A | A | A | A | A | A | A | A | A | A |
| **V4_1_2** | B | B | A | A | A | A | A | A | A | A | A | A | A | A | A | A | A | A | A | A | A |
| **GA26_73_4_1** | B | B | A | A | A | A | A | A | A | A | A | A | A | A | A | A | A | A | A | A | A |
| **d133_42_16_1** | B | A | A | A | A | A | A | A | A | A | A | A | A | A | A | A | A | A | A | A | A |
| **d127_9_21_1** | B | A | A | A | A | A | A | A | A | A | A | A | A | A | A | A | A | A | A | A | A |
| **V4_14_2** | B | A | A | A | A | A | A | A | A | A | A | A | A | A | A | A | A | A | A | A | A |
| **V1_46_3** | A | A | A | A | A | A | A | A | A | A | A | A | A | A | A | A | A | A | A | A | B |
| **d16_76_5_1** | A | A | A | A | A | A | A | A | A | A | A | A | A | A | A | A | A | A | A | B | B |
| **402-18-8-3_2** | A | A | A | A | A | A | A | A | A | A | A | B | B | B | B | B | B | B | B | B | B |
| **371_8_11_4_2** | A | A | A | A | A | A | B | B | B | B | B | B | B | B | B | B | B | B | B | B | B |
| **D14_54_1_2** | A | A | A | A | A | A | B | B | B | B | B | B | B | B | B | B | B | B | B | B | B |
| **GA_18_23_15_2** | A | A | A | A | A | A | B | B | B | B | B | B | B | B | B | B | B | B | B | B | B |
| **V3_70_3** | A | A | A | A | A | B | B | B | B | B | B | B | B | B | B | B | B | B | B | B | B |
| **D13_39_5_2** | A | A | A | A | A | B | B | B | B | B | B | B | B | B | B | B | B | B | B | B | B |
| **GA_17_40_11_2** | A | A | A | A | A | B | B | B | B | B | B | B | B | B | B | B | B | B | B | B | B |
| **d12_149_3_1** | A | A | B | B | B | B | B | B | B | B | B | B | B | B | B | B | B | B | B | B | B |
| **d16_57_13_1** | A | A | B | B | B | B | B | B | B | B | B | B | B | B | B | B | B | B | B | B | B |
| **d12_74_6_1** | A | A | B | B | B | B | B | B | B | B | B | B | B | B | B | B | B | B | B | B | B |
| **GA19_118_5_1** | A | B | B | B | B | B | B | B | B | B | B | B | B | B | B | B | B | B | B | B | B |

**Fig. S2.** Graphical genotype of selected RILs from *YrH52* mapping population. Allele A – resistant allele from *T. dicoccoides* acc. H52; allele B – susceptible allele from *T. durum* cv. LDN. *YrH52* column corresponds to the phenotypes observed at seedling screen with *Pst* at 14 dpi.
